# One-Pot Exosome Proteomics Enabled by a Photocleavable Surfactant

**DOI:** 10.1101/2022.03.18.484933

**Authors:** Kevin M. Buck, David S. Roberts, Timothy J. Aballo, David R. Inman, Song Jin, Suzanne Ponik, Kyle A. Brown, Ying Ge

## Abstract

Exosomes are small extracellular vesicles (EVs) secreted by all cells and found in biological fluids, which can serve as minimally invasive liquid biopsies with high therapeutic and diagnostic potential. Mass spectrometry (MS)-based proteomics is a powerful technique to profile and quantify the protein content of exosomes but the current methods require laborious and time-consuming multi-step sample preparation that significantly limit throughput. Herein, we report a one-pot exosome proteomics method enabled by a photocleavable surfactant, Azo, for rapid and effective exosomal lysis, protein extraction, and digestion. We have applied this method to exosomes derived from isolated mammary fibroblasts and confidently identified 3,466 proteins and quantified 2,288 proteins using reversed-phase liquid chromatography coupled to trapped ion mobility spectrometry (TIMS) quadrupole time-of-flight mass spectrometer. 3,166 (91%) of the identified proteins are annotated in the exosome/EVs databases, ExoCarta and Vesiclepedia, including important exosomal markers, CD63, PDCD6IP, and SDCBP. This method is fast, simple, and highly effective at extracting exosomal proteins with high reproducibility for deep exosomal proteome coverage. We envision this method could be generally applicable for exosome proteomics applications in biomedical research, therapeutic interventions, and clinical diagnostics.

---

Exosomes are nano-sized extracellular vesicles (EVs) of endosomal origin ranging between 30 and 150 nm in diameter and package biomolecular markers reflecting the cells that secrete them.^1^ The exchange of exosomal nucleic acid, metabolite, and protein cargoes via exosome binding and uptake represents an increasingly recognized mechanism of intercellular communication.^2,3^ Recently, exosome-mediated communication has attracted significant attention for its involvement in diseases such as cancer, cardiovascular dysfunction, and neurodegeneration.^4,5^ Since exosomes are secreted by all cells and are present in all biological fluids, they represent attractive targets as minimally-invasive liquid biopsies to diagnose disease, understand disease progression, and serve as therapeutic drug delivery vehicles.^3,6^ Hence, it is important to develop robust techniques for the rapid and reproducible analysis of the biomolecules in exosomes.^7^

Mass spectrometry (MS)-based proteomics is one of the most promising techniques for the global identification and quantification of proteins,^8–10^ and has recently been employed to characterize exosomal protein cargoes.^1,7,11–20^ Typically, bottom-up proteomic methods are used in these MS-based analyses of EVs, but the sample preparation typically involves the use of MS-incompatible detergents for protein extraction, overnight and/or multiple enzymatic digestions, and lengthy multidimensional chromatographic separations.^15–20^ The elongated experimental time and complexity required to improve exosomal proteome coverage in these previously established methods significantly reduce the throughput and reproducibility, limiting the potential of MS-based proteomic analysis of exosomes in translational and clinical applications.

To overcome these limitations, we developed a new method for exosome proteomics with a one-pot preparation of exosomes using a photocleavable surfactant, Azo^21^, to simplify protein extraction and expedite digestion **(Figure 1)**. After Azo-assisted digestion and surfactant photodegradation, the peptides are analyzed using trapped ion mobility spectrometry (TIMS)-quadrupole time-of-flight mass spectrometer (Bruker timsTOF Pro) with parallel accumulation-serial fragmentation (PASEF)^22^ for improved sensitivity and coverage. We have previously shown Azo promoted protein solubilization including both membrane and extracellular matrix proteins, enabled rapid digestion, and yielded reproducible protein identification and quantitation.^21,23,24^ Here, we show that Azo can be used to simultaneously lyse and extract proteins from exosome samples and then assist rapid trypsin digestion. Using this Azo-enabled method, exosome extraction and sample preparation require only ~2.5 h, compared to traditional methods employing overnight digestion and/or lengthy prefractionation that may take 16 to 24 h total.^15–20^ The subsequent LC-TIMS-MS/MS analysis enhanced by PASEF drastically increases sensitivity without increasing analysis time, obviating the need for multidimensional LC or prefractionation, and increasing the total number of proteins identified in the extraction.^24,25^ We have applied this method to analyze mammary fibroblast-derived exosomes. Mammary fibroblast-derived exosomes contribute to the pool of breast tumor exosomes which have previously shown involvement in breast cancer metastatic niche formation and growth^13^. Our one-pot, photocleavable surfactant-assisted sample preparation is simple, rapid, and yields deep exosomal proteome coverage.

**Figure 1.**
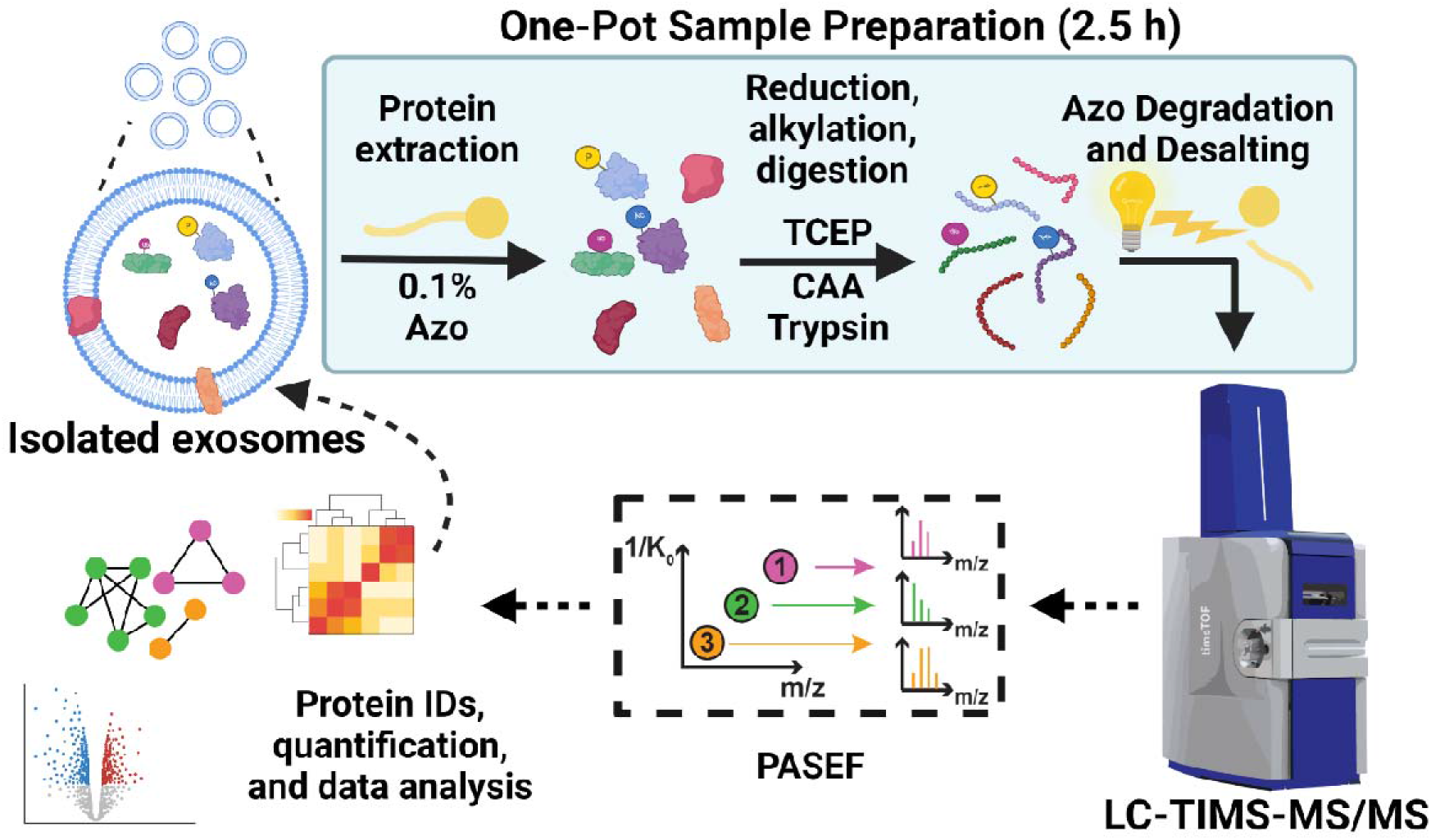
Schematic representation of Azo-based exosome proteomics method. Exosomes are lysed and proteins extracted in 0.1% Azo. Proteins are then reduced and alkylated simultaneously using TCEP and chloroacetamide, followed by Azo-aided rapid trypsin digestion (1 h). The resulting peptides are irradiated with a high-powered UV lamp for 5 min to degrade the Azo surfactant, then extractions are centrifuged and desalted using solid-phase C18 tips before LC-TIMS-MS/MS analysis. PASEF enables deep and robust MS/MS analysis to drastically improve proteome coverage. The resulting data are searched with MSFragger and further analyzed using Perseus.

## Results and Discussion

Exosomes were isolated from mammary fibroblasts by differential ultracentrifugation^26^ and characterized by nanoparticle tracking analysis (NTA) with a Dv50 median particle diameter of 136 nm (50% of the sample exosomes by volume were below 136 nm in diameter) **(Figure 2A)**. This size is consistent with the values of exosome hydrodynamic diameters, as reported previously.^27^ Exosome isolation methods vary across studies and offer differing balances between specificity, yield, and efficiency, though ultracentrifugation is the most common.^28,29^ Isolated exosomes were aliquoted to microcentrifuge tubes and treated with 0.1% Azo in 25 mM ammonium bicarbonate supplemented with Halt Protease Inhibitor Cocktail to extract proteins in triplicate (hereafter referred to as samples 1, 2, and 3) to assess the performance of the method. Following incubation at 37 °C on a thermoshaker for 10 min and then in a bath sonicator for 10 min, 25 mM TCEP (tris(2-carboxyethyl)phosphine) and 50 mM CAA (chloroacetamide), were simultaneously added for a 30 min combined reduction and alkylation step **(Figure S1)**. After pH adjustment to approximately 8.5, trypsin was added and rapid digestion was carried out at 37 °C for 1 h before quenching with neat formic acid (10% v/v). Surfactant degradation was carried out using a 100◻W high-pressure mercury lamp (Nikon housing with Nikon HB-10101AF power supply; Nikon), after which samples were centrifuged and desalted using Pierce C18 tips. Samples were then analyzed in triplicate (200 ng per injection replicate) using a Bruker nanoElute fitted with a C18 column (IonOpticks, 25 cm length, 75 μm inner diameter, 1.6 μm particle size, 120 Å pore size) coupled to a timsTOF Pro operating in data-dependent acquisition (DDA) PASEF mode for MS/MS data collection.

**Figure 2.**
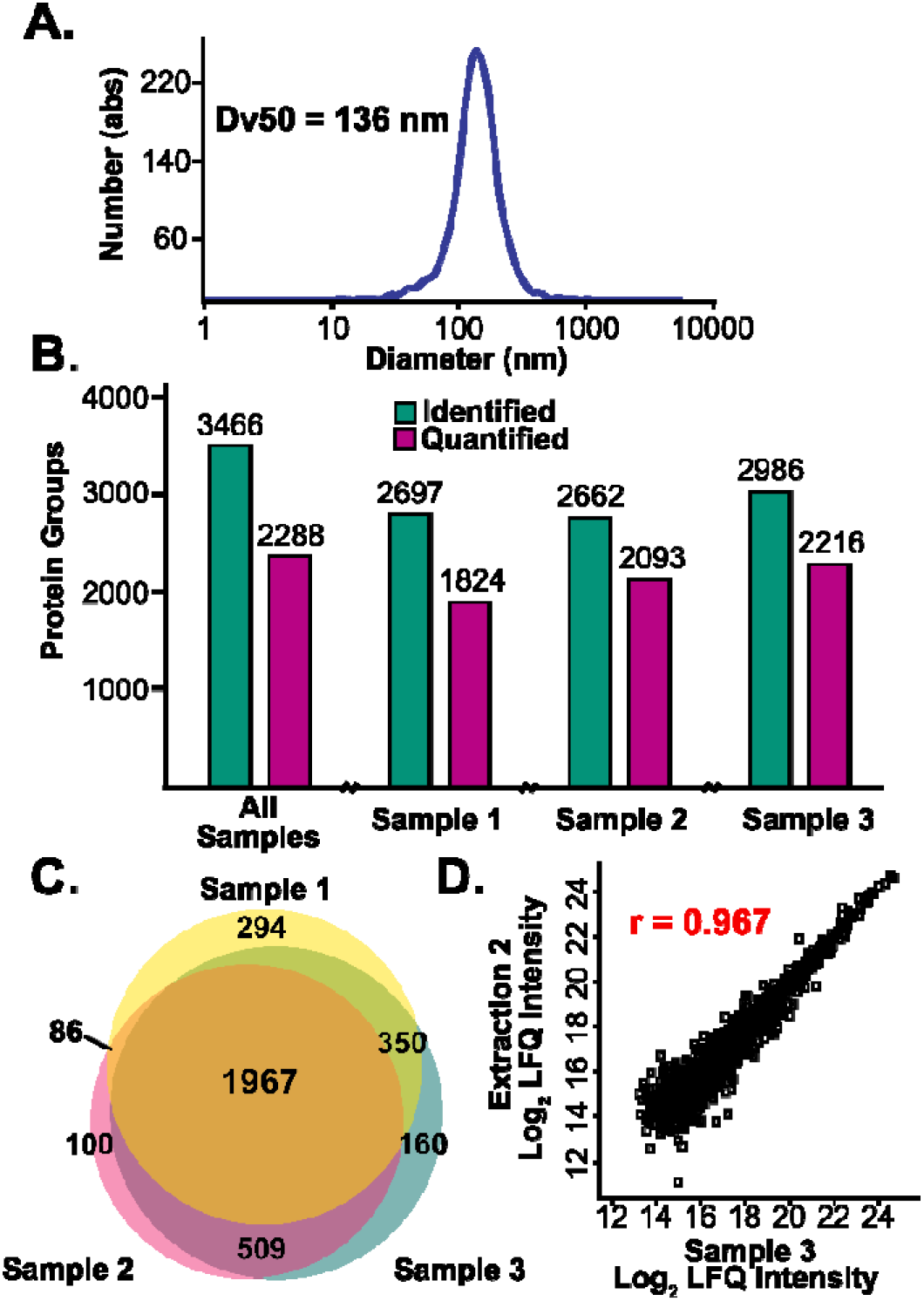
Assessment of exosome isolation, method efficacy, reproducibility, and quantitation in three mammary fibroblast exosome samples. (A) Nanoparticle tracking analysis (NTA) results show a particle size distribution characteristic of exosome samples, with a Dv50 of 136 nm. (B) Bar graph showing protein groups identified (green) with at least one unique peptide spectral match (PSM) and quantified (purple) in at least two samples. Median LFQ intensities from injection replicates of each sample were used for quantitation. From all samples combined, 3,466 protein groups were identified and of those 2,288 were quantified in two of the three extractions. (C) Venn diagram showing the overlap in identified protein groups between samples. We identified 1,967 protein groups common to all three and 2,912 in at least two. (D) Representative scatter plot with associated Pearson correlation coefficient (PCC) depicting a pairwise relationship between log_2_ transformed LFQ intensities of samples 2 and 3.

Identification and label-free quantification (LFQ) following MS analysis were carried out using MSFragger (1% FDR) with IonQuant.^30^ Log_2_ transformed protein LFQ intensities from the three samples showed normally distributed values that spanned similar ranges, suggesting that the extractions were reproducible **(Figure S2 and S3)**. Full lists of identifications can be found in **SI Table 1**.

We identified a total of 3,466 unique protein groups across all samples and quantified 2,288 unique protein groups in at least two of three **(Figure 2B)**. Disaggregated unique protein group counts for each sample and the corresponding number of quantified proteins match roughly with the unique peptides identified (**Figure 2B and S4**). We observed a high degree of overlap in protein group identifications across samples, with 1,967 protein groups being reliably identified in all three **(Figure 2C)**.

To evaluate the quantitative reproducibility of the method, pairwise comparisons of log_2_ transformed LFQ intensities for samples were plotted against each other as scatter plots with associated Pearson correlation coefficients (PCCs).^24,30^ The median LFQ intensity from injection replicates was used from each sample for the downstream analyses. Representative injection replicates for sample 3 showed highly correlated data with an average PCC of r = 0.957, and the total ion chromatogram (TIC) of all three replicates remains consistent in intensity throughout the RPLC-MS analysis **(Figure S5)**. The scatter plots comparing samples show high correlations with PCCs off r = 0.842, r = 0.882, and r = 0.967 with samples 2 and 3 compared in **Figure 2D**. Assuming consistent extraction efficiency across replicates, variability in the PCCs of the samples is likely influenced by their compositions, since sample 1 exhibited the lowest overall number of quantified protein groups, and thus less overlap with the others.

To benchmark the performance of this Azo-enabled method, we compared all the identified protein groups from our samples to those reported in ExoCarta,^31^ a manually-curated online exosome database that collects proteins and nucleic acids identified in previous exosome studies for researchers to use, and Vesiclepedia,^32^ which is similar but includes results from broader categories of EVs including microvesicles (**Figure 3**). 3,166 (91%) protein groups we identified in this study from mammary fibroblast exosome samples are annotated in the ExoCarta and Vesiclepedia databases (**Figure 3A)**. Among them, 2390 protein groups are annotated in both ExoCarta and Vesiclepedia databases, 32 annotated only in ExoCarta, and 744 annotated only in Vesiclepedia. This demonstrates the high quality of our exosome isolation and proteome coverage, since ExoCarta includes data from many cell/tissue types, and we achieve 45% overlap in proteins (2,422) using one cell line, despite the highly heterogeneous nature of exosome content.^6^ In addition, our samples showed generally high abundance based on normalized LFQ intensity of proteins from the ExoCarta top 100 list that collects the most frequently identified protein markers in exosomes **(Figure S6)**.

**Figure 3.**
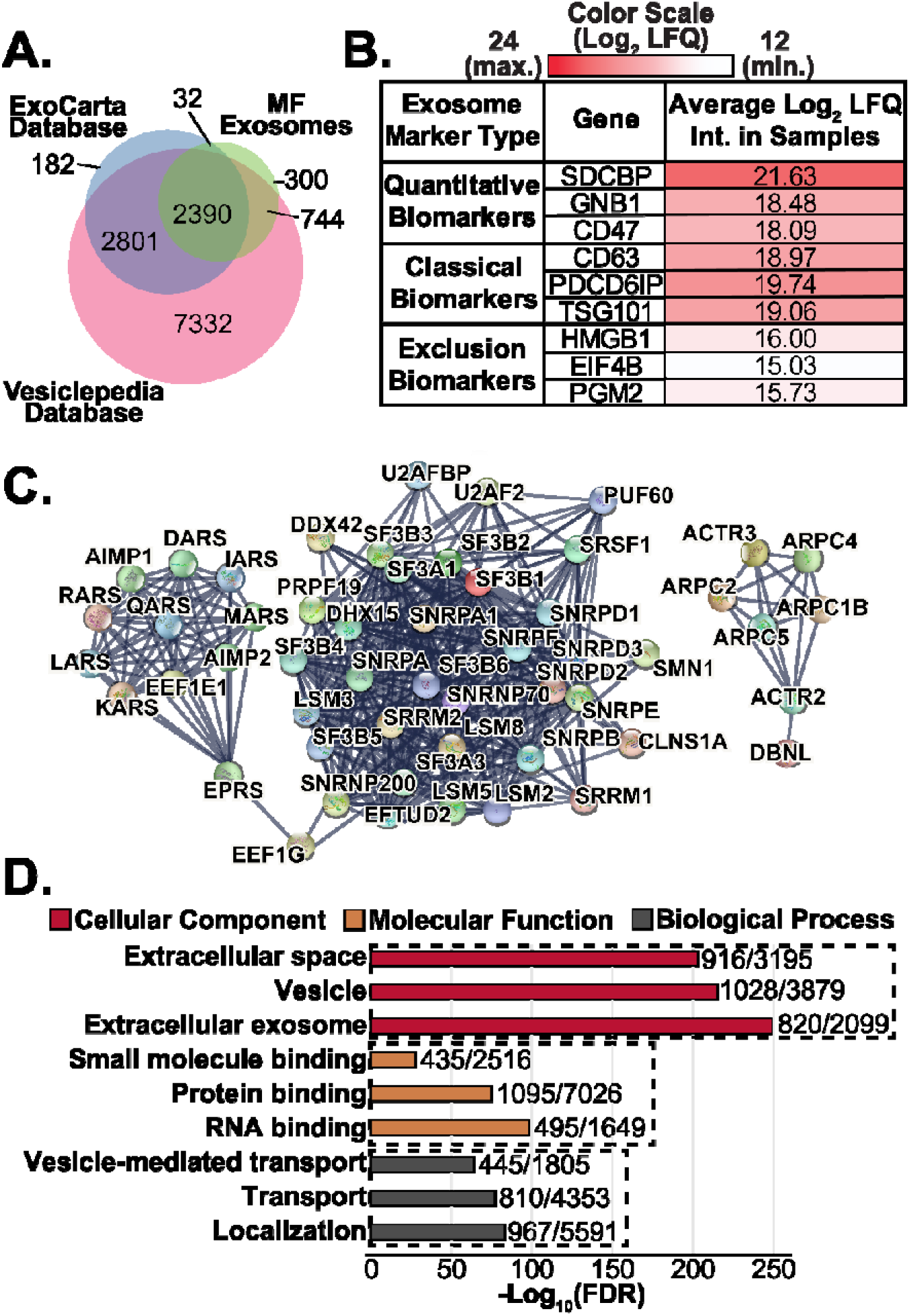
Exosome proteomics results enabled by the Azo surfactant. (A) Venn diagram showing the overlap between proteins identified from mammary fibroblast (MF) exosome samples (3466), ExoCarta database of exosome-specific proteins (5324), and Vesiclepedia database of proteins generally found in EVs (13186). (B) Table showing classical, quantitative, and negative exosomal protein markers, color-scaled to the overall range of observed LFQ intensities of the proteins in mammary fibroblast exosome samples. (C) Zoom-in of STRING PPI network. Clusters show (left to right) aminoacyl-tRNA synthetases, spliceosome proteins, and a group of actin-related proteins. (D) Gene ontology results from STRING PPI analysis, shown grouped by cellular component, molecular function, and biological process. Overall STRING map shows a PPI enrichment p-value of less than 1.0×10^−16^, indicating significant interactions. GO terms are shown with their corresponding −log10 normalized false-discovery rates and the ratio of counts in the network to expected counts.

A small percentage (<9%) of the proteins identified in the Azo-extracted samples were not annotated in ExoCarta or Vesiclepedia. GO analysis of these proteins showed they were largely composed of histone proteins, ATP synthase subunits, and tRNA synthetases which have been previously studied^34^ in exosomes. ATP synthase subunits and histones could potentially be packaged into vesicles or be cellular remnants leftover from isolation. Normalized LFQ intensities of these 300 proteins show their overall low abundance, especially compared to expected markers from ExoCarta and other previous studies (**Figure S6 and S7)**. Given the low abundances of these proteins, it is possible that they are uniquely present in mammary fibroblast exosomes and have yet to be included in the database from previous studies, or are from residual cellular remnants. Additional variation could be attributed to the high degree of heterogeneity in exosomal contents across cell types and exosomes of the same origin.^6,26^

We further investigated exosomal protein markers by plotting the averaged, transformed LFQ intensities of specifically identified characteristic exosome protein markers **(Figure 3B)**. Traditionally, tetraspanins (CD9, CD81, CD63) are used as exosomal markers for proteomics and Western blotting experiments.^1^ However, a recent study of exosome heterogeneity has shown the ubiquity of these markers to be questionable and proposed 22 quantitative markers including SDCBP, GNB1, and CD47 using data from quantitative MS experiments studying exosomes derived from 14 cell lines and isolated by density gradient, size-exclusion chromatography, and ultracentrifugation.^20^ They also proposed 15 “exclusion” markers that were consistently found to be low in abundance including HMGB1, EIF4B, and PGM2. From these proposed quantitative and exclusion markers, we selected three representative proteins to show their respective enrichment or depletion in our samples. We also selected the three traditionally-used markers CD63, PDCD6IP (or ALIX), and TSG101, of which PDCD6IP and TSG101 were included in the list of quantitative markers. The LFQ intensities for the exclusion markers in our samples fell below the lower quartile for all replicates and near the upper quartile for traditional and quantitative inclusion markers **(Figure 3B and S2)**. These numbers are shown in **Figure 3B** with the color scaled to reflect the range of log_2_ LFQ intensities observed from our samples. The relative intensities of these specific positive and negative exosomal markers provide proteomic support for the purity of the exosome isolation.

Using STRING^34^ network analysis, we next assessed the interactions present in proteins identified from the samples **(Figure 3C and S8)**. We plotted the quantified protein groups showing the highest LFQ intensities across the replicates in a protein-protein interaction network that showed an enrichment p-value of < 1.0 x 10^−16^ indicating a significant enrichment. The resulting densely connected network contained clusters of interactors with relevant functional roles in exosomes, including aminoacyl-tRNA synthetases,^35^ splicing factors and other spliceosome-associated proteins,^36^ and actin-related proteins ^37^. Specific proteins from these clusters have been plotted independently in **Figure 3C** but are shown in the full context of the STRING-based PPI network in **Figure S8**.

Further analysis was carried out using the built-in GO capability of STRING to display terms pertaining to the locations, processes, and functions of networked proteins along with their associated FDRs and ratio of gene counts in the network to expected counts **(Figure 3D)**. The most confidently assigned cellular components from the GO analysis were all related to extracellular spaces or vesicles **(Figure S9)**. The majority of confidently-assigned functions and processes pertained to binding and localization respectively. The importance of binding and localization as roles of exosomal protein cargo has already been well-established, as they are fundamentally important to vesicular targeting and uptake.^38^ Additional gene ontology (GO) analysis of the proteins not currently annotated in either ExoCarta or Vesiclepedia is shown in **Figure S10**. Here we have identified the tRNA synthetase cluster, which is consistent with a previous study.^35^ Nevertheless, they are yet to be annotated in ExoCarta or Vesiclepedia database.

To summarize, we developed an Azo-enabled exosome proteomics method capable of one-pot exosome lysis and protein extraction to allow high-throughput sample processing and proteomic analysis. The use of Azo for effective protein extraction and rapid digestion before MS analysis expedites exosome sample preparation. Moreover, the use of TIMS front-end separation and PASEF provides high sensitivity MS/MS analysis for deep proteome coverage. Notably, 3,466 proteins were identified from mammary fibroblast exosome samples and 2,288 proteins were reliably quantified with high reproducibility. This one-pot Azo-enabled exosome method is simple, rapid, and robust, making it amenable for exosome proteomics in general. We envision that this method can help advance the studies of exosomal protein cargoes to understand the role of exosomes in intracellular communication and accelerate the use of exosomes in therapeutic interventions and clinical diagnosis.

## Associated Content

Materials and Methods: Exosome isolation; Sample preparation; Data acquisition; and Data Analysis

SI Figures

Figure S1: Schematic representation of one-pot sample preparation;

Figure S2: Box and whisker plots showing ranges of log_2_ normalized LFQ intensities;

Figure S3: Histogram showing normally-distributed counts of log_2_ transformed LFQ intensities;

Figure S4: Bar graph showing number of unique peptide spectral matches;

Figure S5: Representative injection replicates showing reproducibility of analysis;

Figure S6: ExoCarta top 100 list color-scaled to sample LFQ intensity;

Figure S7: Color-scaled heat map of proteins identified in MF samples and not annotated in databases;

Figure S8: STRING protein-protein interaction (PPI) network map;

Figure S9: Gene ontology results of top 2000 proteins by LFQ intensity;

Figure S10: Gene ontology of proteins identified in MF samples and not annotated in databases SI Tables

Table S1: Protein Output from MSFragger Search

Table S2: Peptide Output from MSFragger Search

Tables S3-S5: Gene ontology results (cellular component, molecular function, and biological process) for top 2000 proteins by LFQ intensity

Tables S6-S8: Gene ontology results (cellular component, molecular function, and biological process) for proteins identified in MF samples and unannotated in databases

## Notes

The authors declare the following competing financial interest(s): The University of Wisconsin−Madison has filed a provisional patent application P180335US01, US serial number 62/682027 (7 June 2018) on the photocleavable surfactant. Y.G., S.J., and K.A.B. are named as inventors on the provisional patent application. The University of Wisconsin−Madison has filed a provisional patent application P220113US01 (1 November 2021) on this work. Y.G., S.J., K.M.B, and K.A.B are named as inventors on the provisional patent application.

## Acknowledgements

This research is supported by NIH R01 GM117058 (to S.J. and Y.G.). Y.G. would like to acknowledge NIH R01 GM125085, R01 HL096971, and S10 OD018475. D.S.R. also acknowledges the support from the American Heart Association Predoctoral Fellowship Grant No. 832615/David S. Roberts/2021. K.A.B. would like to acknowledge the Vascular Surgery Research Training Program Grant T32 HL110853. S.M.P would like to acknowledge NIH R01 CA206458. Graphics in main Figure 1 and Figure S1 were created with clipart from www.BioRender.com.

