## Supporting Information for "One-Pot Exosome Proteomics Enabled by a Photocleavable Surfactant"

### Materials and Methods

**Materials.** All reagents were purchased from Millipore Sigma (St. Louis, MO, USA) and Fisher Scientific (Fair Lawn, NJ, USA) unless noted otherwise. All solutions were prepared with HPLC-grade water (Fisher Scientific). Trypsin Gold was purchased from Promega (Madison, WI, USA). Azo was synthesized in house as described previously.[1]

**Exosome isolation.** Cells were grown to 5 million per plate and cultured in OptiMEM for 48 h, after which conditioned media was collected. Media was centrifuged (CRB Optima Ultracentrifuge, SW32-Ti Rotor) at 300 × *g* for 10 min. to pellet cells, then 2000 × *g* for 20 min and then 10,000 × *g* for 30 min to pellet microvesicles. Vivacell concentrators were washed with 70 mL deionized water at 1000 × *g* for 10 min, and 70 mL of the supernatant from the previous centrifugation was transferred to the filter. Concentrators were centrifuged for 8 min at 1000 × *g*, the resulting flow-through was discarded, and additional supernatant was added to the concentrator. This process was repeated until all of the supernatant was transferred to the concentrator, and the final volume of concentrated material was approximately 12 mL.

12 mL of concentrated material was added to a 14 × 95 mm tube and spun at 100,000 × *g* at 4 °C for 4 h (CRB Optima Ultracentrifuge, SW40 swinging rotor). The supernatant was removed and resuspended in 3 mL PBS. This resuspension was centrifuged at 100,000 × *g* at 4 °C for 2 h in new tubes, and the resulting supernatant was then discarded. Pelleted exosomes were resuspended in 100 μL PBS and flash-frozen in 10 μL aliquots in liquid nitrogen for storage at -80 °C. One aliquot was reserved for the characterization of exosome concentration and diameter using nanoparticle tracking analysis (Particle Metrix Zetaview).

**Sample preparation.** Exosome aliquots were thawed and dissolved in 0.1% working concentration of Azo (4-hexylphenylazosulfonate), 25 mM ammonium bicarbonate, and 1X HALT protease and phosphatase inhibitor cocktail. Aliquots were then placed on a thermoshaker at 37 °C and 600 rpm for 10 min. Samples were placed in a bath sonicator for 10 min and then normalization of protein concentration was performed using the Bradford assay. For reduction and alkylation of disulfide bonds, samples were treated simultaneously with 25 mM TCEP and 50 mM chloroacetamide (CAA)[2] and incubated at 37 °C and 600 rpm on a thermoshaker for 30 min, after which they were treated with 1 M ammonium bicarbonate to adjust pH to the active range for digestion (~8.5). Digestion was performed by treating samples with a 50:1 (w/w) protein:trypsin ratio and incubating them for an hour on a thermoshaker at 37 °C and 600 rpm.

Trypsin digestion was quenched by the addition of small volumes of neat formic acid to reduce sample pH to 2. UV degradation of Azo was then performed using a high-powered mercury lamp to irradiate samples for 10 min, after which they were spun down at 21,000 × *g* for 15 min. To remove degradation products and other contaminating salts, 100 µL Pierce C18 tips were used according to the manufacturer's specifications, and the remaining peptides were resuspended in 0.1% formic acid. Peptide concentrations were determined using absorption at 205 nm from a NanoDrop, using the Scopes method to obtain the extinction coefficient used in the calculations.[3]

**Data Acquisition.** A Bruker timsTOF Pro trapped ion mobility Q-TOF instrument fitted with a captive-spray nano-ESI source and coupled to a nanoElute nanoflow LC was used for all analyses. In each analysis, 200 ng of peptides from exosomal digests was injected onto a C18 column (25 cm length, 75 µm inner diameter, 1.6 µm particle size, 120 Å pore size; IonOpticks). Separations were carried out at 55°C using a stepwise gradient increasing from 2-85% of 0.1% formic acid in acetonitrile and decreasing percent of 0.1% formic acid in water.

To collect MS/MS spectra, the timsTOF Pro was operated in positive mode using DDA-PASEF (data-dependent acquisition parallel accumulation-serial fragmentation) with 10 PASEF MS/MS scans collected over a charge range of 0 to 5. Operating *m/z* was set between 100 and 1700 and a $1/k_0$ range of 0.6 to 1.6 ($V \cdot s/cm^2$) was used with a polygonal mobility filter to exclude singly charged ions. TIMS ramp and accumulation times were set to maintain a 100% duty-cycle, with a ramp time of 100 ms and an accumulation time of 2 ms. Sample injection amounts were normalized by TIC intensity to 200 ng injections of K562 whole-cell lysate.

**Data Analysis.** Identification and protein quantification were performed using MSFragger with IonQuant.[4,5] For MSFragger searches, precursor mass tolerance was set to +/- 20 ppm, and fragment mass tolerance was set to 20 ppm. A maximum of two missed trypsin cleavages were specified and peptide mass was set between 500 and 8,000 Da. For quantitation, default parameters for IonQuant within the FragPipe GUI were used, with match between runs (MBR) enabled. Imported search results were further analyzed in Perseus (ver. 1.1.15.0). After data were filtered for contaminants, values were $Log_2$ transformed, and plots were generated using the resulting LFQ intensities.

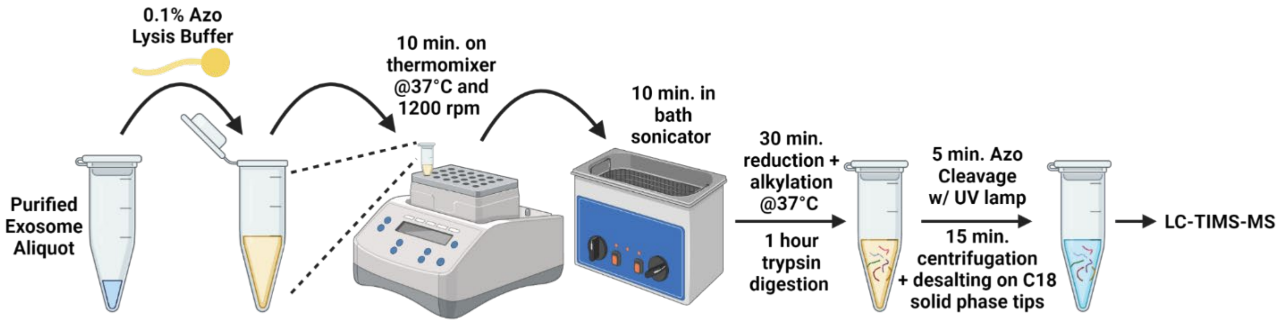


**SI Figure 1.** Schematic representation of one-pot sample preparation for the photocleavable surfactant, Azo-enabled exosome proteomics method. Exosomes are first treated with 0.1% Azo then placed on a thermomixer for 10 min at 37 °C to allow for protein extraction. This is followed by 10 min incubation in a bath sonicator to effectively lyse exosomal membranes. Next, samples are treated with TCEP and CAA for simultaneous reduction and alkylation, while incubating in a 37 °C water bath prior to trypsin digestion (1h). Afterward, the surfactant is degraded by UV and sample cleanup proceeds following centrifugation.

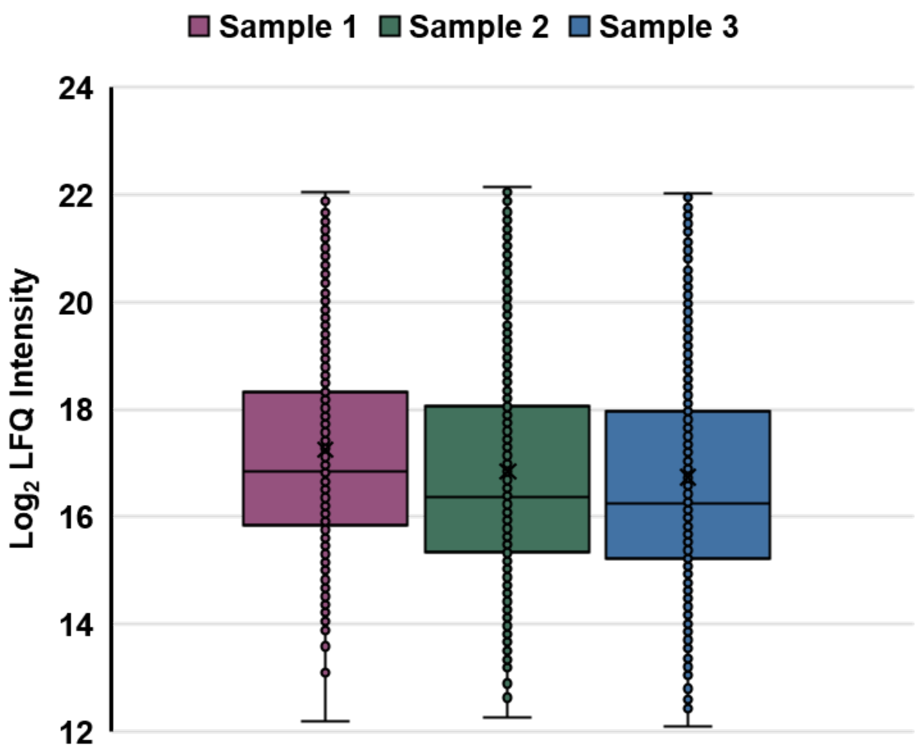


**SI Figure 2.** Box and whisker plots showing ranges of $\log_2$ normalized LFQ intensities, derived from median values from injection replicates for each of the three samples. Outliers are omitted and an exclusive median is used to calculate the ranges shown.

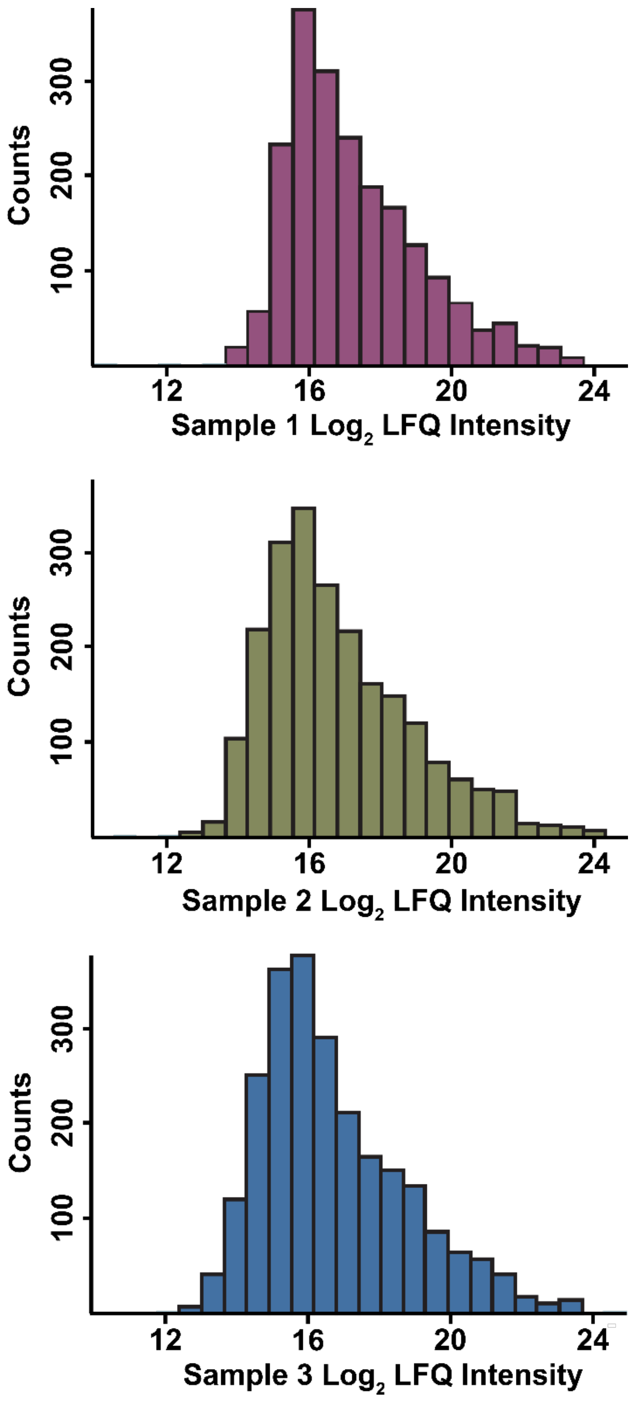


**SI Figure 3.** Histogram showing normally-distributed counts of $\log_2$ transformed LFQ intensities for median values taken from injection replicates of each of the three samples. Results demonstrate reproducible quantification.

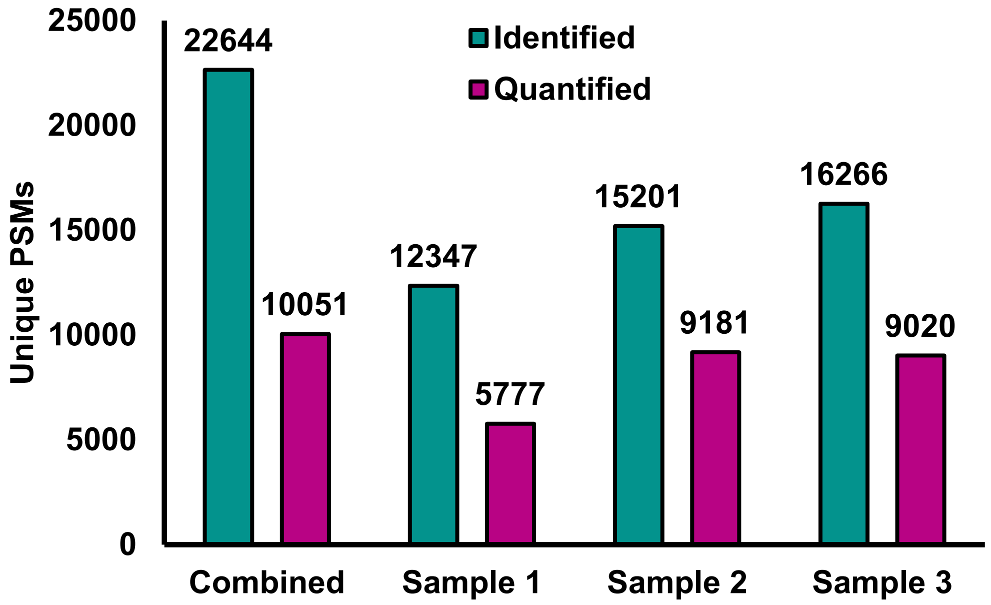


**SI Figure 4.** Bar graph showing the number of unique peptide spectral matches (PSMs) identified in each sample (green) and the number of PSMs with valid LFQ intensities (purple) in two of three samples. For each sample, LFQ intensities were $\log_2$ transformed and the median intensity for each PSM across injection replicates was used.

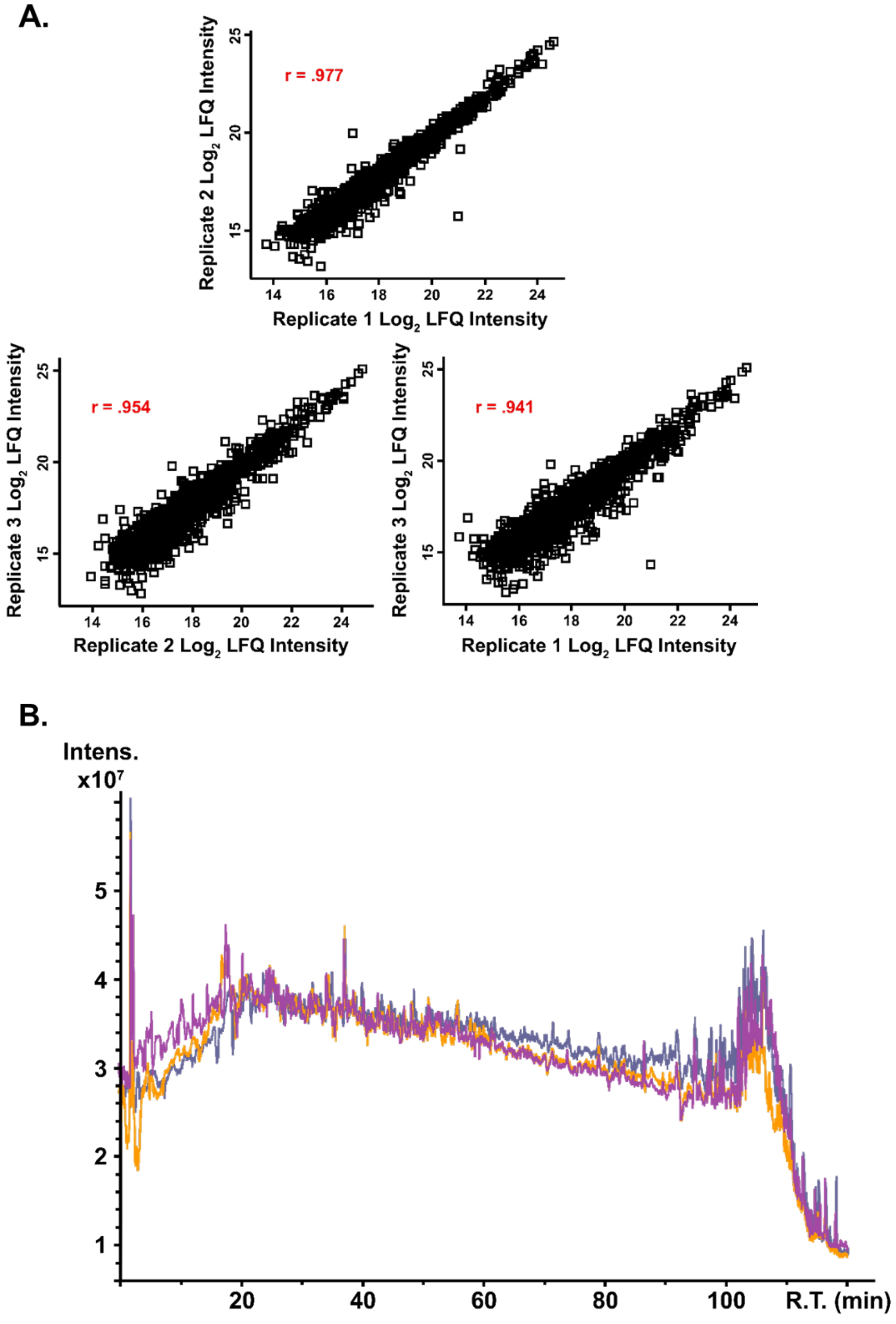


**SI Figure 5.** Representative injection replicates showing reproducibility of analysis. (A) Scatter plots comparing LC injection replicates for sample 3 to each other, with corresponding PCCs. (B) Overlaid total ion chromatogram (TIC) traces from injection replicates of sample 3.

**Color Scale (Log$_2$ LFQ)**

24 (max.) — 12 (min.)

| Rank in Top 100 | Gene Symbol | Rank in Top 100 | Gene Symbol | Rank in Top 100 | Gene Symbol |
|---|---|---|---|---|---|
| 1 | CD9 | 35 | HSPA5 | 68 | ACTG1 |
| 2 | PDCD6IP | 36 | SLC3A2 | 69 | KPNB1 |
| 3 | HSPA8 | 37 | HIST1H4A | 70 | EZR |
| 4 | GAPDH | 38 | GNB2 | 71 | ANXA4 |
| 5 | ACTB | 39 | ATP1A1 | 72 | ACLY |
| 6 | ANXA2 | 40 | YWHAQ | 73 | TUBA1C |
| 7 | CD63 | 41 | FLOT1 | 74 | TFRC |
| 8 | SDCBP | 42 | FLNA | 75 | RAB14 |
| 9 | ENO1 | 43 | CLIC1 | 76 | HIST2H4A |
| 10 | HSP90AA1 | 44 | CCT2 | 77 | GNB1 |
| 11 | TSG101 | 45 | CDC42 | 78 | THBS1 |
| 12 | PKM | 46 | YWHAG | 79 | RAN |
| 13 | LDHA | 47 | A2M | 80 | RAB5A |
| 14 | EEF1A1 | 48 | TUBA1B | 81 | PTGFRN |
| 15 | YWHAZ | 49 | RAC1 | 82 | CCT5 |
| 16 | PGK1 | 50 | LGALS3BP | 83 | CCT3 |
| 17 | EEF2 | 51 | HSPA1A | 84 | AHCY |
| 18 | ALDOA | 52 | GNAI2 | 85 | UBA1 |
| 19 | HSP90AB1 | 53 | ANXA1 | 86 | RAB5B |
| 20 | ANXA5 | 54 | RHOA | 87 | RAB1A |
| 21 | FASN | 55 | MFGE8 | 88 | LAMP2 |
| 22 | YWHAE | 56 | PRDX2 | 89 | ITGA6 |
| 23 | CLTC | 57 | GDI2 | 90 | HIST1H4B |
| 24 | CD81 | 58 | EHD4 | 91 | BSG |
| 25 | ALB | 59 | ACTN4 | 92 | YWHAH |
| 26 | VCP | 60 | YWHAB | 93 | TUBA1A |
| 27 | TPI1 | 61 | RAB7A | 94 | TKT |
| 28 | PPIA | 62 | LDHB | 95 | TCP1 |
| 29 | MSN | 63 | GNAS | 96 | STOM |
| 30 | CFL1 | 64 | RAB5C | 97 | SLC16A1 |
| 31 | PRDX1 | 65 | ARF1 | 98 | RAB8A |
| 32 | PFN1 | 66 | ANXA6 | 99 | MYH9 |
| 33 | RAP1B | 67 | ANXA11 | 100 | MVP |
| 34 | ITGB1 | | | | |

**SI Figure 6.** ExoCarta top 100 list, which collects most frequently identified protein markers in exosomes. Markers are color-scaled to the overall range of observed LFQ intensities in the mammary fibroblast exosome samples. High abundance of these markers shows agreement with previous exosome studies.

**Color Scale**
**($Log_2$ LFQ)**

**24 (max.)** — **12 (min.)**

| | | | | | |
|---|---|---|---|---|---|
| AARS1 | CIAO2B | GPATCH4 | LSM8 | PTPA | SSPN |
| ABRAXAS2 | CIAO3 | GPKOW | LY96 | PYCR3 | SSU72 |
| ACTR6 | CMC2 | GRK2 | MACROH2A1 | QARS1 | SSX1 |
| ADGRF5 | CNPY4 | GSDME | MAD2L1 | QKI | SUMF1 |
| ADPRS | COA4 | GTF2A1 | MARS1 | RAB3IL1 | SUSD6 |
| ADSS2 | COA6 | GTF2E1 | MDFIC | RABEP2 | SYT11 |
| AK5 | COA7 | GTF2F2 | MEAK7 | RABL6 | TACC3 |
| AKAP8L | COMMD8 | GTSF1 | MENT | RACK1 | TARS1 |
| ANKRD40 | COPRS | H1-10 | METTL9 | RAD9A | THUMPD1 |
| ARFGAP2 | COPZ2 | H1-2 | MGAT4B | RARS1 | TIGAR |
| ARL6IP4 | CPSF4 | H1-4 | MIER1 | RECQL5 | TIMM10 |
| ASCC1 | CRYBG1 | H1-5 | MIX23 | RELB | TIMM10B |
| ATP5F1A | CWC15 | H3-2 | MORF4L1 | RELCH | TMEM115 |
| ATP5F1B | CYRIB | H3-3A | MPC2 | RETREG3 | TMEM123 |
| ATP5F1C | CZIB | H3C1 | MRE11 | RFLNB | TMEM184C |
| ATP5F1D | DAP | H3C15 | MRPL24 | RMDN1 | TMEM199 |
| ATP5ME | DARS1 | H4C1 | MRPS10 | RNF24 | TMEM258 |
| ATP5MF | DPH2 | HAX1 | MSR1 | RO60 | TMEM263 |
| ATP5MK | ECPAS | HDGFL2 | MT-ATP6 | RPAP3 | TNS2 |
| ATP5PB | EFCAB14 | HGH1 | MT-CO2 | RTF1 | TOMM70 |
| ATP5PD | EIF4EBP1 | HNRNPLL | MTREX | RTRAF | TOX4 |
| ATP5PF | EIPR1 | HSBP1 | MYG1 | RWDD4 | TRAFD1 |
| ATP5PO | ELOB | HSP90AB4P | NAF1 | S1PR3 | TRIM27 |
| B4GAT1 | ELOC | HYPK | NAGPA | SAMSN1 | TRMT10C |
| BABAM2 | ELP1 | IARS1 | NARF | SAP30BP | TSC22D2 |
| BMPER | EML1 | IGHD | NARS1 | SARS1 | TSC22D4 |
| BPNT2 | ENDOG | IGHV1-18 | NAXE | SELENOF | TVP23C |
| C11orf98 | ENSA | IGKV3-7 | NECTIN2 | SELENOP | TXN2 |
| C18orf25 | EPRS1 | IGKV3D-20 | NECTIN3 | SEMA4B | TXNL4A |
| C4orf3 | ERBIN | IGLV1-47 | NIBAN1 | SENP3 | UBE2L5 |
| CA8 | ERI3 | JMJD6 | NIBAN2 | SEPTIN10 | UFD1 |
| CACNA1I | ERO1A | JPT1 | NOL10 | SEPTIN11 | UMAD1 |
| CALHM5 | ESF1 | JPT2 | NRDC | SEPTIN2 | UNC50 |
| CAMTA2 | FAM20B | KARS1 | NUFIP2 | SEPTIN7 | URI1 |
| CARS1 | FAM234A | KCT2 | OCC1 | SEPTIN8 | VARS1 |
| CAVIN1 | FBXL15 | KIAA1143 | OGA | SEPTIN9 | VPS9D1 |
| CAVIN2 | FBXO30 | KIAA1549L | OLFML1 | SGCZ | WARS1 |
| CAVIN3 | FKRP | KIFBP | ORC4 | SHTN1 | WASHC2A |
| CCDC177 | FLRT1 | KIRREL1 | PBDC1 | SKA3 | WASHC3 |
| CCDC43 | FOXG1 | KPNA7 | PCYT1A | SLC38A7 | WASHC4 |
| CCN1 | GARS1 | KRI1 | PIP4P1 | SLC49A4 | WASHC5 |
| CCN2 | GASK1B | KTI12 | PIP4P2 | SLC66A1 | WDR13 |
| CCN3 | GATA1 | KYAT3 | PLAC9 | SLX9 | WFDC12 |
| CDC42EP4 | GCN1 | L1RE1 | PLPP3 | SMAP | WNT2 |
| CDKN2AIP | GEMIN6 | L3HYPDH | POLR3D | SMNDC1 | YARS1 |
| CEMIP2 | GET3 | LAMTOR5 | PPP1R11 | SNU13 | ZDHHC8 |
| CERT1 | GFUS | LARS1 | PQBP1 | SNX11 | ZNF775 |
| CFAP298 | GINM1 | LDAH | PRRT1 | SQOR | ZNF806 |
| CHPF | GLMP | LNPK | PRSS3P2 | SRPRA | ZNRD2 |
| CHTOP | GOLM2 | LRCH2 | PSME3IP1 | SS18 | ZPR1 |

**SI Figure 7.** Heat map of proteins not annotated in ExoCarta or Vesiclepedia but identified in our mammary fibroblast exosome samples. Markers are color-scaled to the overall range of observed LFQ intensities.

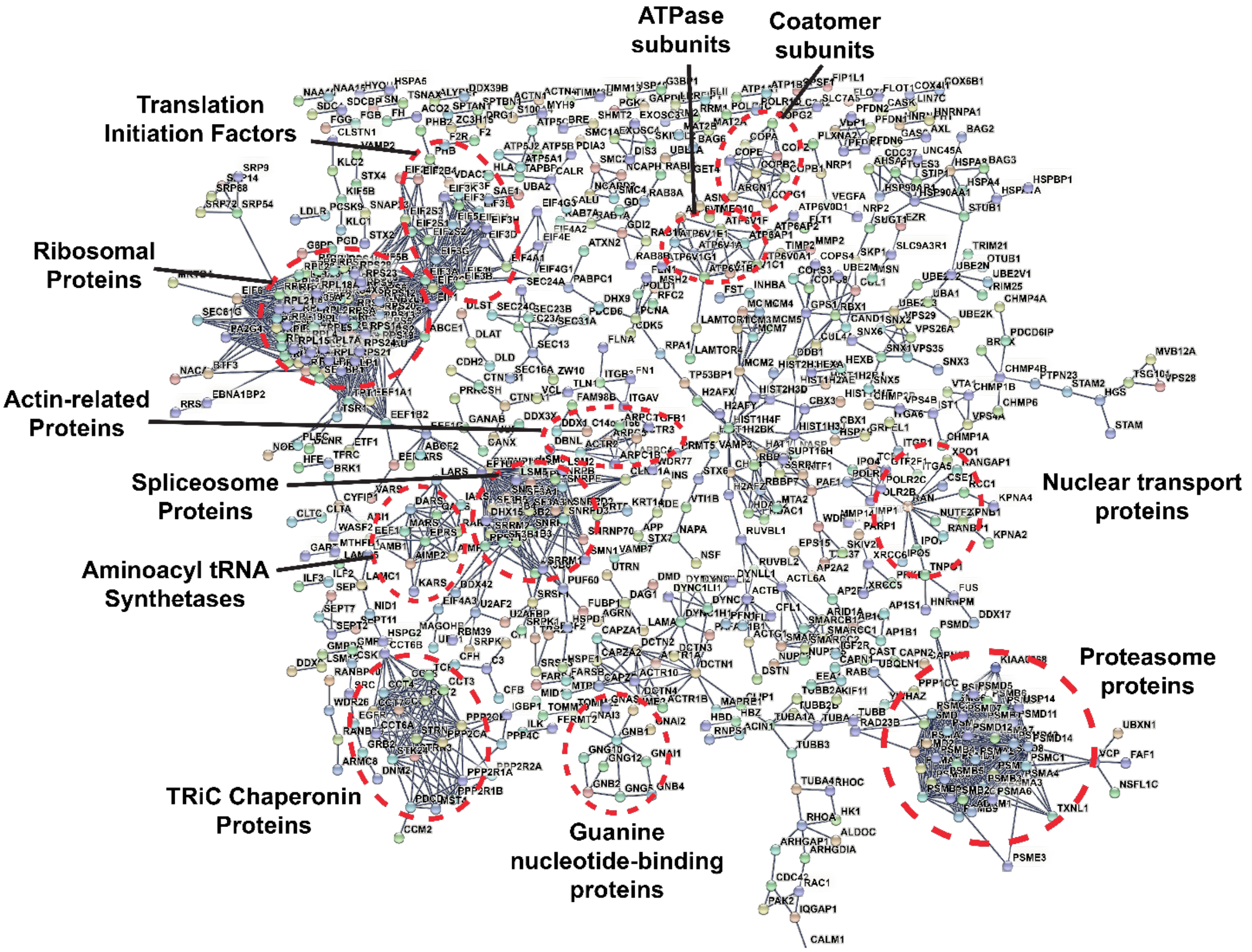


**SI Figure 8.** STRING protein-protein interaction (PPI) network map. Network has a PPI enrichment p-value of < 1e-16, indicating a significant biological connection as a group. A confidence level of 0.9 and the experimental evidence type from STRING were used for the shown interactors. Notable clusters of interactors corresponding to protein complexes or pathways are circled in red and labeled.

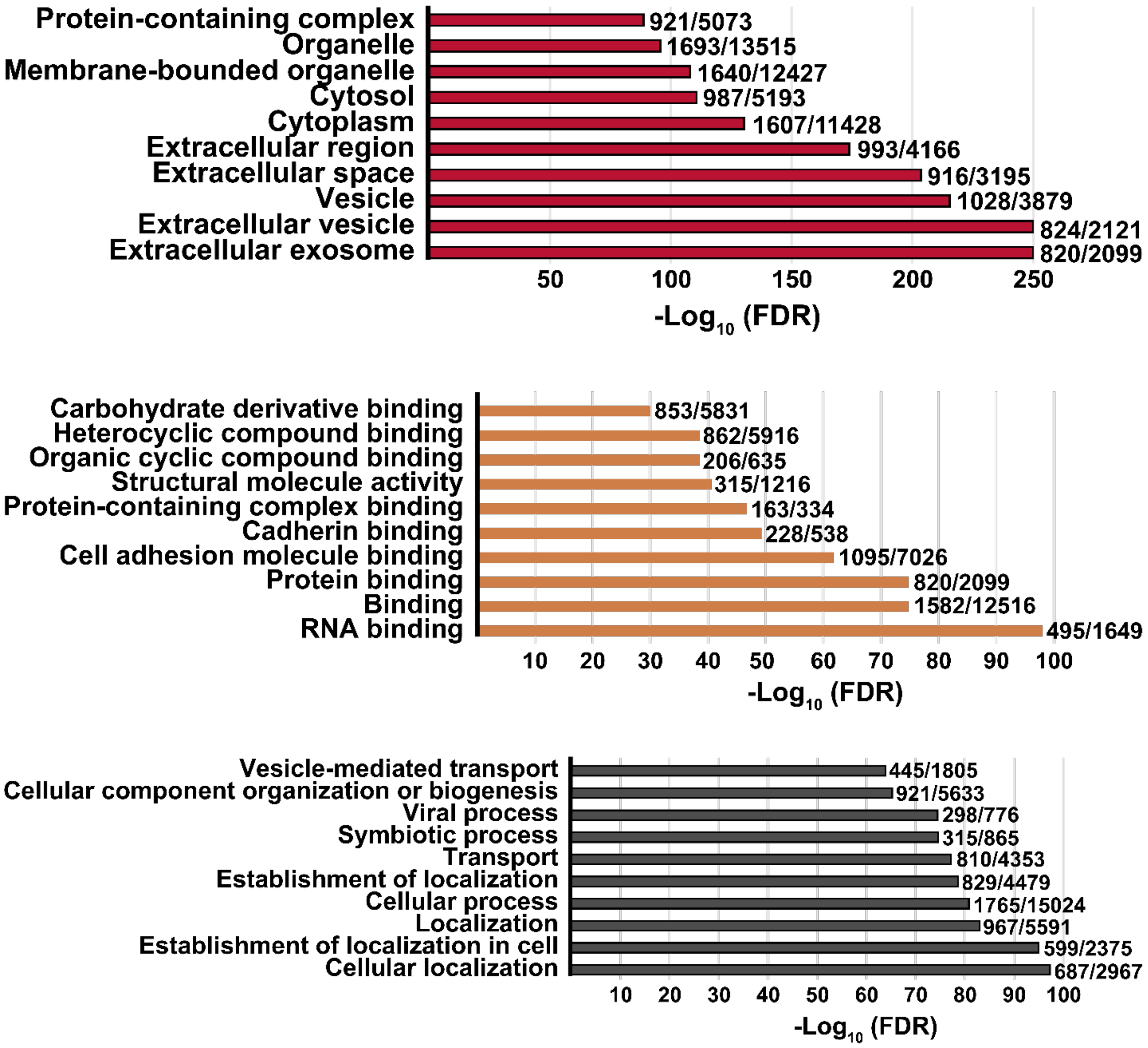


**SI Figure 9.** Results from GO analysis of the top 2000 protein groups by LFQ intensity, showing terms for cellular component (red), molecular function (orange), and biological process (grey) with associated $-\log_{10}$(FDR) values and ratios of gene counts in network to background genes for the top ten highest $-\log_{10}$(FDR) terms in each category.

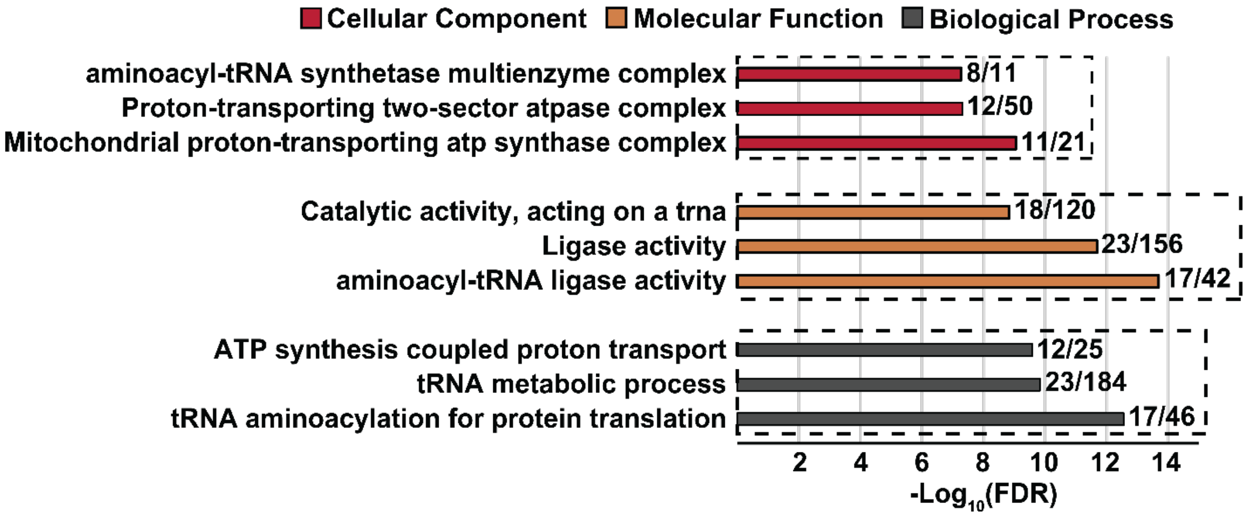


**SI Figure 10.** GO analysis from STRING network of proteins identified in the mammary fibroblast-derived exosomes samples using our method which were not found in ExoCarta or Vesiclepedia databases. Plot shows top three GO cellular component, molecular function, and biological process terms by their $-\log_{10}$ (FDR) and the ratio of observed to background genes in the network. Of the 300 proteins identified in this study which are not annotated in either database, primary categories included ATP synthase subunits, tRNA ligases, and histones.
